# Validation of a Radio frequency identification system for tracking location of laying hens in a quasi-commercial aviary system

**DOI:** 10.1101/2023.02.16.528820

**Authors:** Sabine G. Gebhardt-Henrich, Alexander Kashev, Matthew B. Petelle, Michael J. Toscano

## Abstract

Cage-free housing is increasingly chosen in Europe, North America, and Australia as an animal-welfare friendly farm system for laying hens. However, hens are kept in large numbers in those systems which makes checking for health and welfare difficult and individuals cannot be identified. Tracking systems like radio frequency identification allow researchers to monitor these individuals almost continuously. Individual tracking data has revealed substantial individual variation in movement patterns, however, in recent studies, only a subset of animals per flock was tracked. We applied an RFID tracking system to monitor all 1125 laying hens of a flock, which were divided into 5 pens of 225 birds each in a barn with an aviary system. In each pen, 26 antennas were placed on the edges of three tiers and in the litter. For validation purposes, 3 hens in 2 connected pens were fitted with colored backpacks. They were recorded on video and their location throughout the pen was taken from the video and compared with registrations from the RFID system. For 93% of compared transitions, the RFID data matched the observational data regarding the tier or litter whereas the value fell to 39% for specific antennae. When the antennae on the litter were excluded for the validation, the match on tier-level was at least 98% but on antenna-level it remained lower than 50%. The sensitivity of the detection of tiers/litter but not antennae differed among the three hens. We conclude that the RFID tracking system was suitable for studying the movement pattern of individual hens among tiers in an aviary system in a reliable way but tracking birds on the litter needs to be improved.

## Introduction

Cage-free housing systems for laying hens may contain tens of thousands of animals. Although considered welfare-friendly, cage-free housing systems including aviaries are known to entail risks concerning health (e.g. parasites, infections) and animal welfare (e.g. damaging behaviours like feather-pecking and cannibalism) (Platz, et al., 2009; Blatchford, et al., 2015; Louton, et al., 2017; Li, et al., 2019; Ali, et al., 2020). In principle, aviaries are designed to offer essential functional areas to the hens like aerial perches for (nighttime) roosting, secluded nest areas for laying, and a litter area for exploratory behavior and dust-bathing. However, individual birds access these areas to a different extent (Rufener, et al., 2018) which is known to correlate to various health risks (Rufener, et al., 2019; Ali, et al., 2020).

Tracking individuals in large groups of identically looking laying hens is a challenge that can either be attempted by visually marking the animals or by an electronic tracking system (for reviews see Li et al., 2020; Neethirajan (2022). Visually tracking hens in a three-dimensional aviary system where birds can move to places where they are invisible due to equipment or conspecifics is difficult, time consuming, and limited. Various technologies including Infrared (Rufener, et al., 2018), Radio Frequency Identification (RFID) (Zhang, et al., 2016; Sibanda, et al., 2019), and other (reviews by Siegford et al., 2016; Brown-Brandl et al., 2019) types of systems have been successfully used to track individuals within the aviary in order to measure individual movement patterns and the amount of time spent in the functional areas. Despite these efforts, tracking is typically limited to a subset of the flock or for a limtied time which might not suffice in certain research projects, e.g. heritability estimates for breeding programs. In any case, all automated tracking devices should be validated with video observations (Iserbyt et al., 2018) using the instances when hens are tracked and visible. Therefore, the aim of this study was to validate an RFID system with the capacity to track a much larger number of individual laying hens in an aviary by assessing the accuracy of registrations. For this purpose, we tracked three focal animals within a larger flock of 450 hens within a commercial aviary.

## Methods

### Ethical note

The use of animals was approved by the Veterinary Office of the Kanton of Bern (BE136/2020) on 10-FEB-2021 and met all Cantonal and Federal regulations for the use of animals in scientific research.

### Barn-setup and RFID system

Twenty six 12-field SPEED antennae (length: 75 cm) of a passive 125 kHz RFID System (Gantner Pigeon Systems GmbH, Schruns, Austria) were placed at different locations in a Bolegg Terrace aviary system (Vencomatic Group, Eersel, NL) (Fig. 1). The antennae were encased in plastic, connected to reading devices which were connected by multiplexers (Moxa, New Taipei City 242, Taiwan) to a computer. A similar system was described in Gebhardt-Henrich et al. (2014). On each tier at each side of the aviary structure (upper, nestbox, lower) as well as in the litter, three antennae were put side-by-side joining at the short end. Additional antennae were placed on each side of the wintergarden although not evaluated in this effort. As a test trial for future experiments on a large number of birds, all birds in 5 pens of a barn with 20 pens with 225 birds per pen were fitted with a glass tag (HITAGS 4×22mm, 125KHz, HTS256) in a custom-developed leg band (Fig. 2). If a tag was detected by an antenna a time stamp and the identities of tag and antenna were written into a .csv file every 0.1 s. However, if a tag remained on the same antenna for a 10 s period, the registration was not repeated in order to limit the size of the generated files. The maximal vertical reading distance of all antennae was about 15 cm and the horizontal reading distance was close to 0 cm. Three hens in a pen that was connected at the level of the litter to a neighboring pen for free movement between the two pens wore color-coded back-packs that were visible on video recordings. One observer watched videos recorded between April 21^st^ and 29^th^, 2021 until a total of 10 hours of video were scored, on which at least one hen with a custom-made backpack (Fig. 3) for identification was visible. Based on the recorded video (30 fps), the location of those hens walking, standing, or sitting on the antennae and the pen at each change of location with the respective video time stamps was entered into a spreadsheet. Additionally, the observer noted whether the identification of the hen was certain or uncertain due to poor visibility.

**Figure 1:**
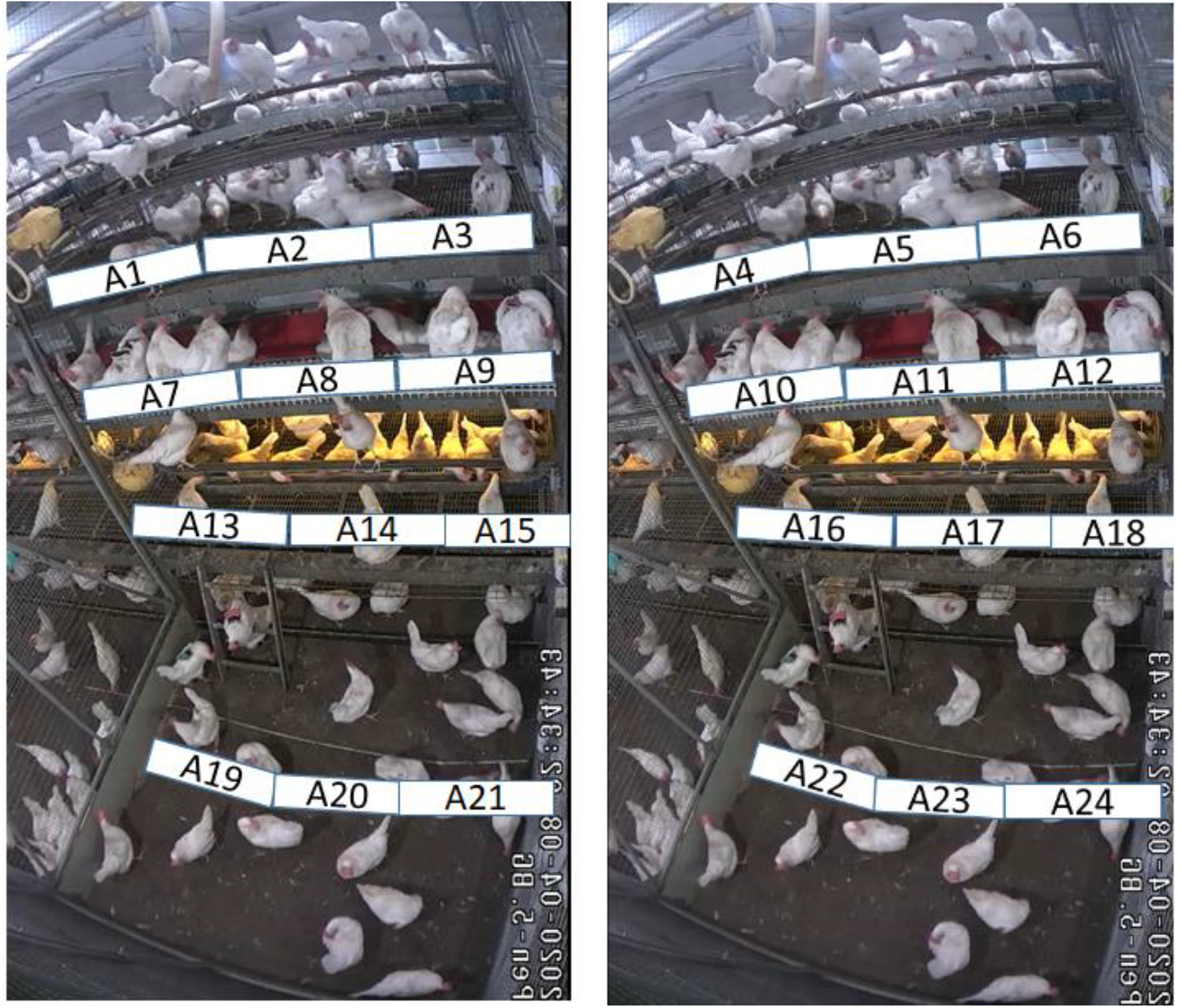
View of both sides of the aviary with the location of 24 of the 26 antennae. Two antennae were located in front and behind a pophole leading to a wintergarden available on one side of the aviary only (left). These 2 antennas were not used in the validation trial.

**Figure 2:**
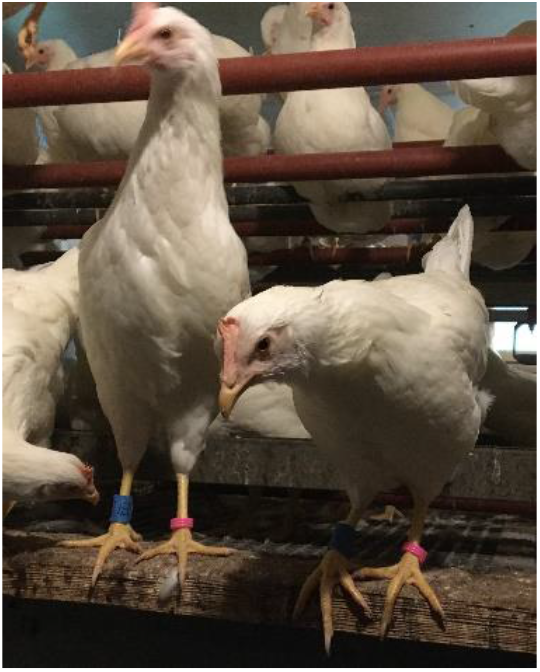
The blue legbands on the right legs contain the RFID tag.

**Fig. 3.**
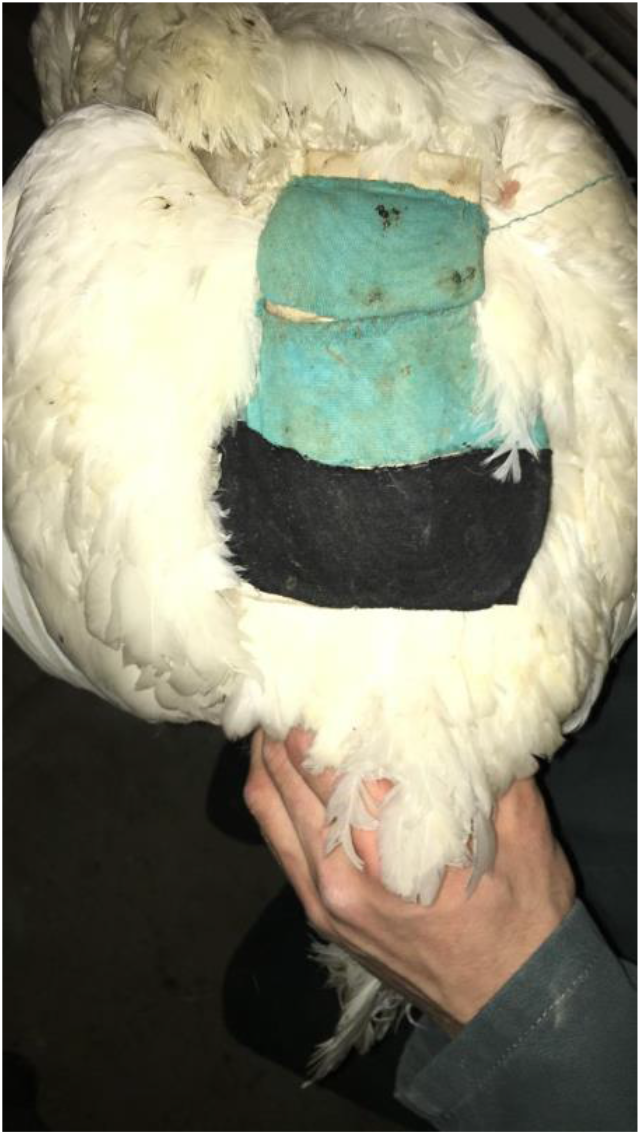
The back of a hen wearing a backpack.

### Analyses

Initial data processing of the registrations of the RFID data (date and time stamp to the closest tenth of a second, ID of the bird, ID of the antenna) were done in R (version 4.2.0). For each observation from the video it was noted whether the RFID system had recorded the bird on the same antenna, tier, and side of the aviary, and in the correct pen. In particular, several variables were extracted for the closest RFID registrations in time that matched the hen (see Table 1), and tests were performed to compare the RFID data and coded observations (see Table 2).

**Table 1:** Variables that were extracted from the RFID data for events from coded video observations.

| Variable | Meaning |
| --- | --- |
| Time difference of the closest event | Time difference [s] between the closest (by time) RFID registration for the observed hen and the observation by the observer. |
| Closest antenna | The antenna code recorded in the closest RFID event as stated above. |
| Closest tier | The tier corresponding to the closest RFID event as stated above. |
| Closest side | The side of the aviary corresponding to closest RFID event as stated above. |
| Closest pen | The pen recorded in the closest RFID event as stated above. |
| Time difference of the exact antenna match | Time difference [s] between the closest (by time) RFID registration that matches the antenna and the observation by the observer. |
| Time difference of the tier + side + pen match | Time difference [s] between the closest (by time) RFID registration that matches the triple (tier, side, pen) and the observation by the observer. |
| Time difference of the tier match | Time difference [s] between the closest (by time) RFID registration that matches the tier and the observation by the observer. |

**Table 2:** Categories of agreement between coded video observation events and corresponding RFID data.

| Variable | Meaning |
| --- | --- |
| Side correction needed | The side as recorded by the observer had to be corrected by a second person because there was an obvious error in coding by the observer (antenna coding for the wrong side was used, based on the antenna and side mismatch). This did not involve RFID data. |
| Closest antenna matches | The antenna code recorded in the RFID event for the observed hen with closest timestamp to the coded video observation time is the same as observed. |
| Same antenna within 1 min. | Same as above but the RFID data matches the observed antenna within a 1 min. window. It is not necessarily the match closest in time. |
| Closest tier + side + pen match | The tier recorded by the observer matches the tier (and side) of the antenna code of the RFID registration closest in time for the respective bird. |
| Same tier + side + pen within 1 min. | Same as above but the observer matches the tier (and side) within a 1 min. cutoff. It is not necessarily the match closest in time. |
| Closest pen matches | The pen recorded by the observer matches the pen of the antenna code of the RFID registration closest in time for the respective bird. |
| Closest side of the aviary matches | The side of the aviary recorded by the observer matches the side of the aviary of the antenna code of the RFID registration closest in time for the respective bird. |

The results were entered in a confusion matrix to calculate the sensitivity of the RFID system (true positive cases / sum of true positive and false negative cases) and the time differences between the time stamp of the video and the time stamp of the RFID system were analyzed using *PROC FREQ* and *PROC UNIVARIATE*, (SAS Institute Inc., 2016).

## Results

From the video files, 304 locations of the three birds were detected of which the observer was certain (75.6% of all sightings of birds on antennae). Of these, in 91 % of the cases, the correct tier, side, and pen of the aviary was detected within 1 min. by the RFID system (Tab. 3a). In all but 7 cases, this was also the closest RFID detection in time. In 1 case, the correct tier, but at the opposite side of the aviary was indicated by the RFID system. The correct tier regardless of the side of the aviary and the pen was detected in 93% of the cases. Sensitivity fell precipitously to 39% when the focus was detecting the correct antenna within one minute. In 3% of the cases a wrong pen was indicated and in 2% the wrong side of the aviary.

When the registrations of birds on the antennae situated on the litter were excluded, detection was much better (Table 3b). All sensitivities on tier-level were between 98 and 99% whereas the sensitivities regarding the correct antenna within tier remained below 50%.

**Table 3:** Sensitivities of the detection of locations of birds as seen on the video file by the RFID system. In some cases, the same antenna or tier was registered on RFID as the observer indicated within 1 min. but there was an earlier RFID registration event of another antenna/tier (named closest antenna etc. in time). a) all observations, N = 304. b) registrations on the litter excluded, N = 158.

| a) |  |  |  |
| --- | --- | --- | --- |
| Registered by RFID | N of event = true | Sensitivity | Difference between hens |
| Closest tier and side and pen in time | 271 | 0.89 | P = 0.002 |
| Same tier and side and pen within 1 min. | 278 | 0.91 | P = 0.001 |
| Closest tier and side in time | 279 | 0.92 | P = 0.02 |
| Closest tier in time | 284 | 0.93 | P = 0.0005 |
| Closest antenna in time | 74 | 0.24 | P = 0.12 |
| Same antenna within 1 min. | 120 | 0.39 | P = 0.11 |
| Closest pen in time | 294 | 0.97 | P = 0.02 |
| Same (aviary) side | 299 | 0.98 | P = 0.53 |
| Observer correct | 234 | 0.77 | P = 0.72 |
| b) |  |  |  |
| Registered by RFID | N of event = true | Sensitivity | Difference between hens |
| Closest tier and side and pen | 154 | 0.98 | P = 0.58 |
| Same tier and side and pen within 1 min. | 156 | 0.99 | P = 1.00 |
| Closest tier and side in time | 154 | 0.98 | P = 0.58 |
| Closest tier in time | 157 | 0.99 | P = 0.15 |
| Closest antenna in time | 48 | 0.29 | P = 0.93 |
| Same antenna within 1 min. | 75 | 0.48 | P = 0.0004 |
| Closest pen in time | 158 | 1.00 | N/A |
| Same (aviary) side | 155 | 0.98 | P = 1.00 |
| Observer correct | 156 | 0.99 | P = 0.30 |

The registration of the RFID system was on average 1.6 s (Stderror = 1.9 s) earlier than the video time stamp if the tier identified by the RFID and observer matched and 3.6 s. (2.5 s.) earlier if the antennas by the RFID and observer matched. Neither time differences were significantly different from zero (same tier: Student’s t = 0.82, P = 0.41, N = 293, same antenna: Student’s t = 1.42, P = 0.16, N = 135).

Of the three hens, each accounted for 43.4% (132), 34.9 (106), and 21.7% (66) of all registrations. The hens differed in the sensitivity of the registrations relative to tiers including the litter but not antennae when all tiers included the litter were analyzed. However, with the exclusion of the antennae on the litter, hens only differed when the same antenna within 1 min. was considered. The difference was due to the two birds with the fewer registrations. Of those, one hen had about 5 times more correct than incorrect registrations of the antenna within 1 min. and the other bird had twice as many incorrect than correct registrations of the antenna within 1 min.

## Discussion

The detection rate of birds on the different tiers and in the litter of an aviary system was very high and comparable to other efforts using different RFID systems in poultry with either equal or greater sensitivities (In broilers: Li et al., 2019 (Ultra-high frequency); van der Sluis et al., 2020 (Ultra-Wide Band), laying hens: Sales et al., 2015 (134.2 kHz); Wang et al., 2019; Sibanda et al., 2020 (UHF (915 MHZ))). The findings were also comparable to efforts using non-RFID systems (see review by Siegford et al., 2016) including those in the same barn applying the same ‘zone’ approach (Rufener et al., 2018; Candelotto et al., 2022)but lower than the reliability of 99% of the active low-frequency tracking system by Montalcini et al.(2022). Although overall sensitivity was high, the correct antenna was detected in less than 50% of the cases. The poor detection can be explained by the fact that the antennae were positioned adjacent to each other so that a tag likely could be read intermittingly by both antennae when the hen sat on both. The problem of birds in between antennae has also been a problem for other efforts (van der Sluis et al., 2020). In addition to this problem ‘within’ pens, the problem could also persist ‘across’ pens. As pens were adjacent, antennae of one pen also touched antennae of the neighbouring pen leading to registrations in the ‘wrong’ pen. In case that pens are connected and the movement of birds between pens is studied, this likely error would need to be addressed. For instance, to resolve the problem of false pen registrations, the edges of antennae at the extreme sides of the pen can be physically blocked (Ringgenberg, et al., 2015). In either case, our efforts suggest the benefits of such a validation to help improve accuracy and determine potential solutions. More critically, our results also indicate that the present set-up did not yield adequate precision to tell where across the 225 cm wide tier the hen was located, i.e. we achieved only the registration of the tier and side with acceptable levels. Given our validation results, tracking individuals at the side/tier level is possible, but a higher resolution may be necessary depending on the research question.

In 20 instances, the RFID registration did not match the correct tier. In all but 1 of these cases, the bird was seen on the litter but the antenna immediately above the litter on the first tier was recorded instead. In one mismatch, the hen was seen on the antenna on the highest tier and it was recorded on that tier but on the other side of the aviary. In each of these cases, the hen likely moved faster than the registration window, e.g. up to the first tier / down to the litter or underneath the aviary to the opposite side. Speed of registration has been shown to be a problem with fast moving laying hens with a similar RFID system (Gebhardt-Henrich, et al., 2014). For the current validation, an improved system with faster registration was used. However, it is possible that very fast moving hens may still be missed. The resolution of the timestamp in the csv file generated by the RFID system was 0.1 s. Since it is impossible to synchronize the video system with the RFID system with this accuracy, the time difference between the RFID registrations and the video time stamps are not surprising.

Tier-specific, incorrect registrations also likely result from the set-up of the aviary and the spatial configuration of the antennae. Interestingly, we found almost no mistakes in terms of tier recordings except in the litter. The decreased sensitivity of the litter is likely because birds can more easily enter the area without coming into contact with an antenna. In contrast, a bird transitioning between the upper and nest box tiers would have to step onto an antennae at the edge of each zone. As a solution to improve sensitivity in the litter, we have doubled the number of antennas there with a later setup.

The three hens differed in the sensitivity of the registrations of the tier and the positions where they were observed in the aviary. One hen was mostly seen on the litter while another on the uppermost tier. The hen with the lowest sensitivity scores had fewer registrations but was seen both on the litter and the uppermost tier. The sample size of three hens is too low to draw any conclusions whether certain individuals would differ in the sensitivity of the registration of tiers. However, it is feasible that such a difference exists due to variations in an individual’s behavior (e.g., flying or jumping across antennas) or preference of certain locations in the aviary which are less reliably registered on the antennas. In our dataset the difference in the sensitivities likely resulted from differences in litter use because sensitivities on tier-level no longer differed among hens when registrations of antennae on the litter were excluded. Differences in the registration of the antennae within one min. were due to the 2 hens with fewer registrations and the cause is unknown.

This validation was done before the start of the full experiment so we do not have tracking data from other hens for this period. However, density of hens and equipment of the pen was the same as in the following studies except the addition of a second row of antennas on the litter in following experiments.

It is important to note that a gold standard to determine the positions of the hens does not exist. The observations from the videos were error prone and in almost one quarter of observations the combination of antennas and side of the aviary were impossible and had to be corrected. Mistakes while coding videos occur like in other easy tasks that do not require a high level of conscious attention esp. when the observer is disrupted (Morrison, 2021). In addition, the antennae on the tiers of the aviary could be clearly seen on the videos whereas the exact positions of antennae on the litter were less obvious because they were covered by litter. This could have added to the lower sensitivities of detection on these antennas. Furthermore, it was difficult to synchronize our video and RFID systems with the resolution of less than 1 s. because both systems were not connected to the same network.

In conclusion, the employed RFID system reliably detected the position of hens on the different tiers in an aviary in a reliable way but tracking birds on the litter needs to be improved.

## Acknowledgements

We thank Masha Marincek to observe the hens on the videos. Abdelsatar Abdul Rahman installed and serviced the RFID system daily. Numerous helpers were involved in catching and banding birds.

## Data, scripts, code, and supplementary information availability

Scripts and code are available online on OSF: https://doi.org/10.17605/OSF.IO/UHTSW.

